# Blind deconvolution in autocorrelation inversion for multi-view light sheet microscopy

**DOI:** 10.1101/2021.10.05.463163

**Authors:** Elena Corbetta, Alessia Candeo, Andrea Bassi, Daniele Ancora

## Abstract

Combining the information coming from multi-view acquisitions is a problem of great interest in light-sheet microscopy. Aligning the views and increasing the resolution of their fusion can be challenging, especially if the setup is not fully calibrated. Here, we tackle these issues by proposing a new reconstruction method based on autocorrelation inversion that avoids alignment procedures. On top of this, we add a blind deconvolution step to improve the resolution of the final reconstruction. Our method permits us to achieve inherently aligned, highly resolved reconstructions while, at the same time, estimating the unknown point-spread function of the system.

## Introduction

Image formation in microscopy can be described mathematically by the convolution of the perfectly resolved object with the microscope’s point spread function (PSF) [1]. The latter represents the impulse response of the system and, ideally, should be as narrow as possible to obtain the best image resolution. When the PSF diverges from the ideal delta function, image-restoration techniques may be used to improve the three-dimensional image quality. Among others, deconvolution (the process of inverting the convolution) aims to retrieve a sharper estimation of the object, reducing optical blur and noise by reassigning the energy distribution of the signal according to the PSF.

In Light Sheet Fluorescence Microscopy (LSFM) [2],[3], the illumination and detection optical paths are independent and orthogonal to each other: the PSF depends on both the illumination and detection optics and, typically, results in a function that is more elongated along the detection axis. For this reason, multi-view acquisitions are frequently adopted as a method to obtain isotropic resolution [4],[5]. Moreover, in the presence of slightly absorbing diffusive specimens [6], multi-view acquisition facilitates the reconstruction of the entire imaged volume by accessing favourable views of the object. However, pre-processing steps are necessary to align and fuse all the acquisitions. In a typical multi-view LSFM pipeline, stacks of images are acquired from various angles, they are rotated, aligned, and combined together. Each of these steps requires dedicated care. The alignment procedure involves finding the rigid shift that guarantees the best overlaps of each view against each other. In general, the fusion can be performed by computing simple averages or with more advanced methods, e.g., by considering Poisson statistics [4] or optimized Bayesian strategies [7],[8],[9].

To complicate the reconstruction further, the knowledge of the PSF is not always accessible [10]. In a light sheet microscope, the acquisition of the PSF is performed by acquiring beads placed in the imaging chamber. However, this procedure may be impractical: this is the case in some clearing media [11] where it is difficult to prepare beads phantoms, or in fluidic-based systems [12]. In these cases, blind deconvolution is the only possible approach to increase imaging resolution [10].

To solve the issues of alignment and fusion, we have proposed the Anchor Update (AU) algorithm [13]. The protocol works out a deconvolved reconstruction from the autocorrelation estimation, exploiting the fact that working in the shift space guarantees implicit alignment. This implies that the solution of the autocorrelation problem always returns reconstructions that are aligned intrinsically within sub-pixel resolution [14].

In the present manuscript, we describe a strategy for tomographic reconstruction in multi-view acquisitions corrupted by unknown blurring, that can be utilized in those cases when the PSF cannot be easily measured. To this end, we design a blind deconvolution approach that acts in the autocorrelation space. This permits the formation of inherently aligned reconstructions, increasing the resolution simultaneously without any prior knowledge of the PSF of the system. We take inspiration from the Richardson-Lucy deconvolution (RLD) [15],[16] and its blind implementation [17], extending the blind approach to the autocorrelation domain. First, we introduce the computational methods by validating the protocol thanks to the reconstruction of synthetic samples. Here, generated images for the object and PSF are used as ground truth to assess the convergence of our algorithm. Once validated, we test our reconstruction method on real data acquired by a custom LSFM setup, with the aim of retrieving both the PSF and the aligned reconstruction. Lastly, we discuss the potential of our method in the field of image processing for optical microscopy applications.

## Results and discussion

### Working principle of blind deconvolution

The goal of any blind deconvolution algorithm is to obtain a deblurred version of the original image without any knowledge of the response of the optical system used for image acquisition. It requires solving the deconvolution problem, *o*_*μ*_ = *o* ∗ *h*, recovering *o*, the original object, from the blurred measurement *o*_*μ*_ without knowing the blurring function *h*. There are infinite couples of images and PSFs that may produce the observed blurred output; thus, it may appear as a seemingly impossible problem to solve. It has been shown, however, that meaningful solutions are achievable by having weak, or even null, knowledge of the object under study and of the PSF [10].

When the PSF is unknown, one of the most effective strategies is provided by a modified blind RLD method, which acts alternatively in the object and PSF domain. This protocol, introduced by Fish et al. [6], consists in dividing the deconvolution process into two steps, in which the object and the PSF commute their roles. First, it executes a certain number of RLD iterations to improve the object reconstruction by using a fixed guess for the PSF. Then, the PSF estimation is improved by using the previously reconstructed object as the kernel of another RLD. This cycle is repeated several times until reaching optimal reconstructions.

Since we are interested in aligned multi-view measurements, we propose an extension of this blind method to the autocorrelation space. This approach is aimed at treating alignment and blind deconvolution together. As previously demonstrated by Ancora et al. [14] in the case of known blurring, the inversion of *χ*_*μ*_ via the AU algorithm permits obtaining an aligned and deconvolved reconstruction *o*_*rec*_. In this case, the problem that we try to solve is the inversion of the average autocorrelation estimated from the measured views:

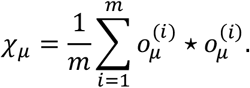

In the previous equation, we have denoted the autocorrelation operator with the symbol ⋆ and with *m* the total number of acquired views. In practice, the problem that we tackle is separating the object from the PSF given the average autocorrelation *χ*_*μ*_ = (*o*_*rec*_ ⋆ *o*_*rec*_) ∗ *H*, where *H* = *h* ⋆ *h* is the autocorrelation of the PSF. In this representation, *o*_*rec*_ is the average of the aligned and deconvolved views *o*^(*i*)^, and *h* represents the PSF that blurs the fused multi-view data, computed as the average of the PSFs of each view. In a multi-view setting, the PSFs simply differ by a rotation angle. Here, we aim to accomplish the same result without having any information about the microscope’s PSF, *h*.

We address the problem as schematized in Fig. 1. We start from the autocorrelation of the measurement *χ*_*μ*_, and we provide two initial guesses: one for the object, *o*_0_, and another for the PSF, *h*_0_. We begin by executing the AU algorithm [13] to recover the object from the blurred autocorrelation as if we knew the PSF. Then, we reverse the role of object and PSF inside the iterative process, and we focus on reconstructing the PSF while keeping the object reconstruction fixed. This cycle is repeated several times until valid reconstructions for both elements are obtained. From now on, we will refer to this strategy as the Blind-AU reconstruction protocol. By denoting with the indexes *t* and *t*′ the iterations within the AU cycles, and with the subscript *k* the outer steps, formally, the Blind-AU can be written as:

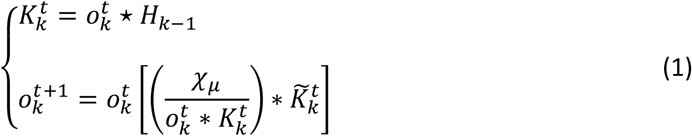

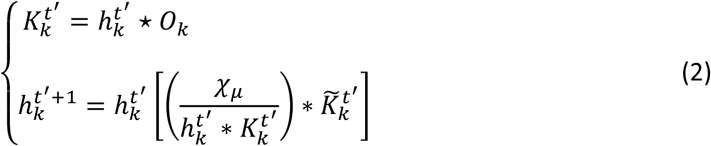

**Figure 1.**
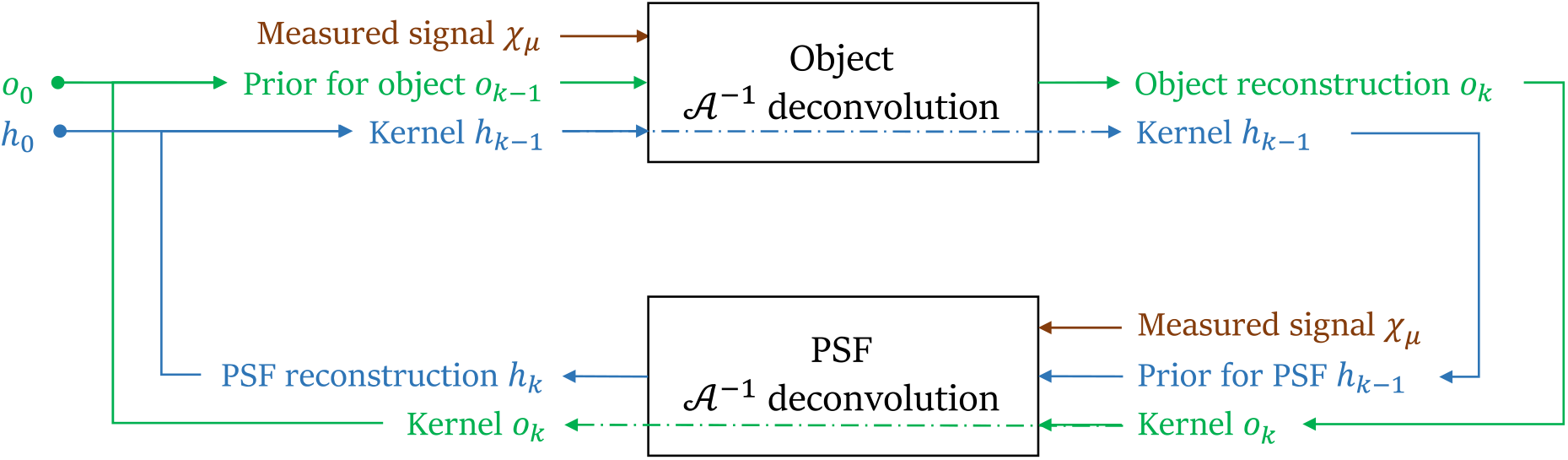
Schematics of blind AU algorithm. The algorithm starts giving as input two initial guesses, one for the object and the other for the PSF. In the first step, autocorrelated signal χ_μ_ is deconvolved and de-autocorrelated to improve the object estimation o_k_. After that, the same signal is used to restore the PSF, maintaining o_k_ unchanged. Upon completion of the outer cycle indicated by arrows, the estimations o_k_ and h_k_ are retrieved, and the process iterated. We choose the number of iterations for each step and the number of times the outer cycle is performed depending on the experimental or simulated conditions.

As previously mentioned, the object and the PSF are denoted with *o* and *h*, whereas we indicate their autocorrelation with the uppercase letters *O* and *H*. Here, the tilde specifies that the element is expressed with reversed coordinates: 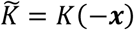.

The code for blind deconvolution is available on GitHub (at the link: https://github.com/danieleancora/blind_AnchorUpdate).

In the following section, we test our new Blind-AU method to reconstruct a synthetic sample, mimicking a misaligned acquisition with differently orientated point spread functions.

### Blind PSF and object reconstructions of synthetic data

In multi-view light-sheet microscopy, the sample is observed from different angles and always exhibits a point-spread function elongated along the detection axis [4]. In Fig. 2, we simulate a typical tomographic section of a sample measured in an LSFM setup. For our case study, we consider a generic sample made by a random arrangement of vessel-like structures, blurred by two PSFs elongated along the vertical (Fig. 2a) and horizontal (Fig. 2b) directions. These directions would correspond to the direction of the mechanical scanning of the sample through the light sheet.

**Figure 2.**
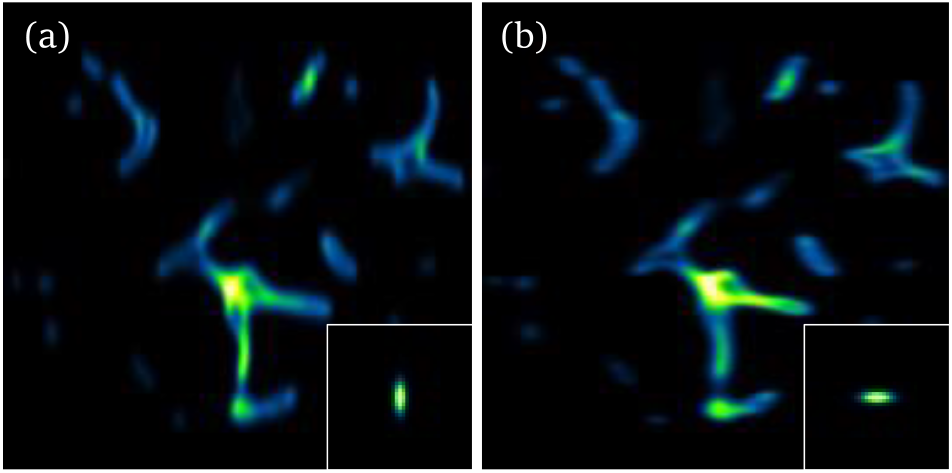
Synthetic sample for blind deconvolution. Blurred and noisy orthogonal views generated starting from an image simulating blood vessels. (a) Synthetic measurement of the ground truth image corrupted by the PSF represented in the inset. (b) Orthogonal measurement with its corresponding PSF.

We test our image reconstruction method with this dataset, aiming at obtaining a tomographic slice sharper than the mere average of both images. The original section of the synthetic sample, which is our ground truth, is shown in Fig. 3a. Aligning and fusing the orthogonal views leads to the blurred reconstruction in Fig. 3b (top-left triangle). The alignment was accomplished by locating the peak of their cross-correlation and compensating for its opposite shift [18]. As a result of the fusion, the blurred reconstruction is affected by the combination of the two orthogonal PSFs that generate a star-like point spread function (Fig. 3c).

**Figure 3.**
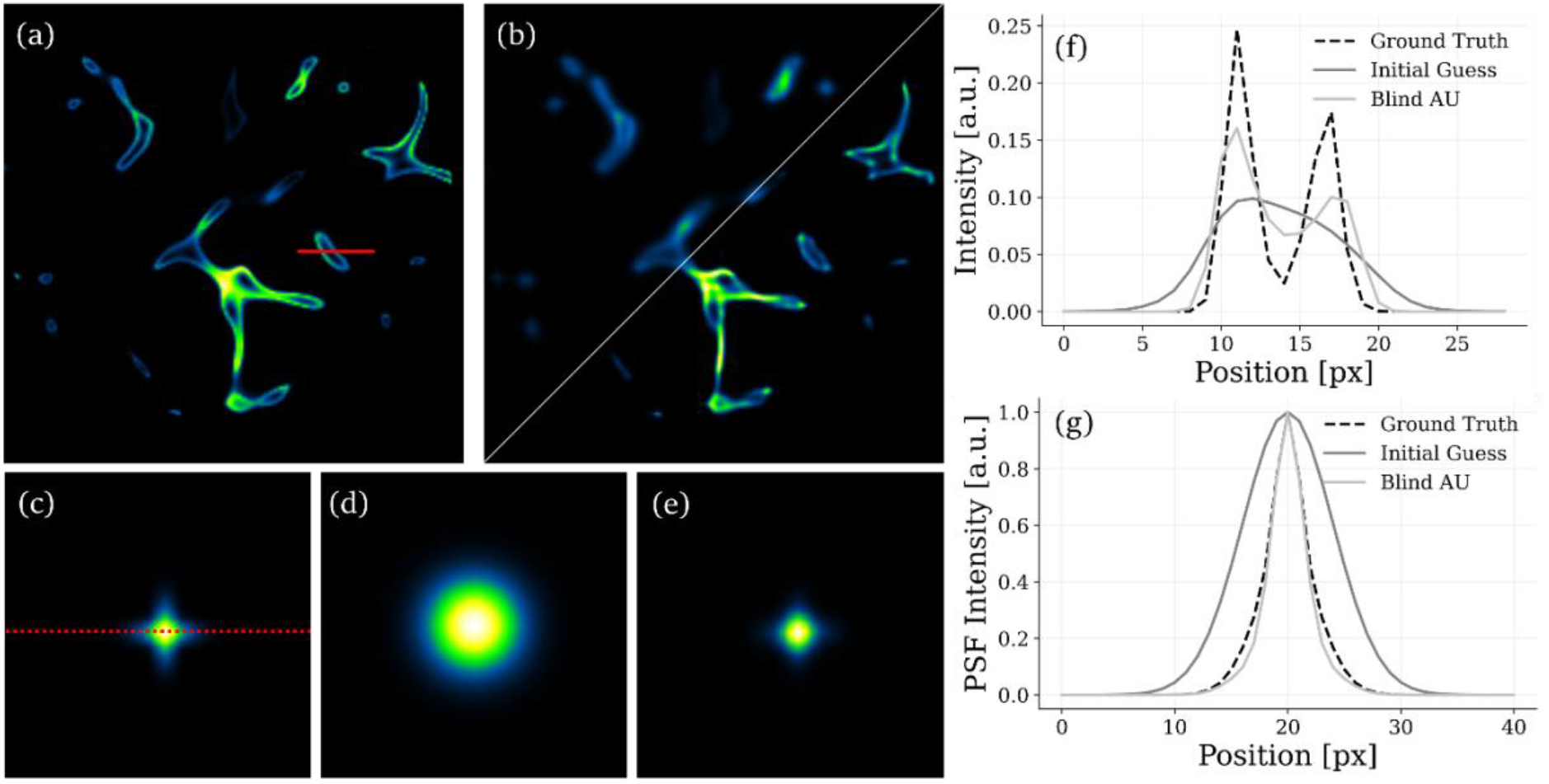
Simulated Blind-AU reconstruction. (a) Original synthetic sample resembling a vessel network. (b) Upper-left triangle: blurred reconstruction of synthetic data obtained by averaging orthogonal views of the object. Bottom-right triangle: reconstruction obtained with our Blind-AU strategy. (c) Blurring kernel resulting from the fusion of orthogonal views. (d) Initial guess for the blurring kernel from which we start our blind reconstruction. (e) PSF kernel obtained at the end of the Blind-AU algorithm. (f) Intensity profiles along the red line drawn in panel (a) of the ground truth, simulated measurement, and Blind-AU result. (g) PSF profiles along the red dashed line drawn in panel c: blurring kernel computed as the average of the PSFs from multiple angles, initial guess for the PSF, and PSF reconstructed at the end of the Blind-AU. That software to generate this figure is available on GitHub (https://github.com/danieleancora/blind_AnchorUpdate).

Our Blind-AU method allows for the reconstruction of both the object, sharper than each single view, and the PSF. Since our algorithm assumes that we do not have any information about the system, we set the initial guess of the PSF to be an isotropic Gaussian with *σ* = 4 *px* (Fig. 3d). This initialization is larger than the ground truth PSF in both directions. On the other hand, we keep the aligned average as the initial guess for the object. Ideally, our protocol will adjust the guess of the PSF to the actual PSF that blurred the synthetic measurements.

We run a total of 2 · 10^5^ AU iterations to obtain the reconstructions: the protocols are executed in steps of 50 iterations for both the object and the PSF, and the outer blind cycle is repeated 2000 times. With this choice, the object and the PSF are processed for 10^5^ iterations each. The result of the object reconstruction is shown in the bottom-right triangle of Fig. 3b. The reconstructed image exhibits higher contrast and enhanced void regions, even when not directly discernible in the blurred version. We use intensity profiles to quantify the resolution increase by comparing the ground truth, the initial guess, and the de-autocorrelated and deconvolved image (Fig. 3f). We observe the profiles along the red line drawn in Fig. 3a. Starting from a hump, in which the opposite walls are indistinguishable, the two peaks of the original image are recovered. The deconvolution process transfers the signal energy from the center to the two opposite lobes. Moreover, the recovered PSF reaches a remarkable result: if we start from an isotropic Gaussian function, the kernel quickly converges towards a star-like shape (Fig. 3g). For completeness, we report that the cross takes form after a few hundred iterations, and its overall shape continues to refine during the following steps.

### Multi-view blind deconvolution of zebrafish samples

After validating the method with synthetic samples, we tested it experimentally in a multi-view light-sheet microscope. We report the reconstruction results obtained with zebrafish (*Danio rerio*) embryos having the vasculature stained with a fluorescent protein. Experimental details are provided in the methods. For this experiment, we acquired four orthogonal views of the specimen, focusing on the central region of the tail. In experimental measurements, we do not have access to the ground truth reconstruction of the specimen, so we compare our reconstructions with the one provided by standard RLD by assuming known the blurring PSF. To obtain this reconstruction, we initially align each acquisition against the other by locating the position of the maximum within their cross-correlation.

Once aligned, we fuse the volumes by calculating their average. Figure 4a shows the maximum intensity projection (MIP) of the fused volume (upper panel) and a tomographic slice along the dashed blue line (bottom panel). We estimate the PSF of a single view by measuring fluorescent nanobeads, fitted with a three-dimensional Gaussian function. Finally, the PSF that blurs the multi-view reconstruction is obtained by rotating the single-view PSFs accordingly to the acquisition angle of each dataset. We use the average PSF to enhance the resolution of the dataset via RLD; the resulting volume is shown in Fig. 4b. For the Blind-AU, we compute the average autocorrelation of each view, as described in the previous section. We recall that this step does not require any alignment process. Then, we feed this estimation to the algorithm described in Eqs. 1–2 and schematized in Fig. 1. After running a total of 25 · 10^4^ iterations, we obtain the result displayed in Fig. 4c. Our reconstruction is sharper and highly resolved compared to the standard fused data and to the deconvolved one (Fig. 4a,b). In particular, we can appreciate the improvement along the tomographic plane (bottom panels of Fig. 4), where our method neatly discriminates the two vessels located in opposite regions of the spine. To quantitatively assess the improvement, we examine the profiles of the three volumes along the direction indicated by the red line in Fig.4a. The plot of the profiles is shown in Fig. 4d. As expected, the RLD gives results sharper than the initial guess. Our method further surpasses this result by better separating the opposite vessels, reconstructing details sharper than standard approaches.

**Figure 4.**
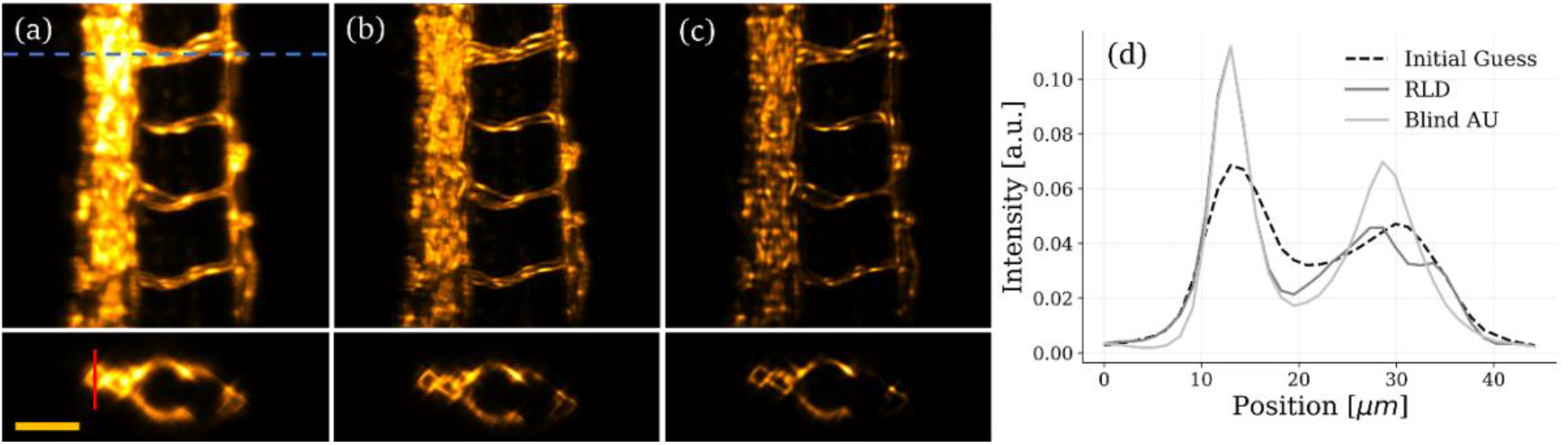
Four-view RL and Blind-AU reconstructions of the tail of a zebrafish embryo. (a) Top: maximum intensity projection (MIP) of the fusion of four orthogonal views (45°, 135°, 225°, 315°) of the zebrafish tail section, used as the initial guess for RLD and Blind-AU algorithms. Bottom: tomographic section of the same sample taken at the location of the dashed blue line. Scale-bar: 50 μm. (b) MIP and tomographic section of the RLD reconstruction. (c) MIP and tomographic section of the reconstruction using the Blind-AU algorithm. (d) Intensity profile along the red line traced on the tomographic section (a) plotted for the initial guess, RL, and Blind-AU reconstruction.

Having assessed the image reconstruction ability, we now focus on the quality of the PSF obtained at the end of the process. We take a slice along the tomographic plane obtained by fusing orthogonal views (Fig. 5a). We first average the autocorrelations of all the views (Fig. 5a). The Blind-AU is initialized with the fused object reconstruction (Fig. 5b) and a completely random guess for the PSF. The object reconstruction and PSF are optimized alternatively with our method. After running 5 · 10^4^ iterations, we obtain the reconstructions of the object displayed in Fig. 5c, and of the PSF in Fig.5d. Also, in this case, the increase in the resolution can be verified by the intensity plot along the red line of panel a: our method reconstructs edges that are sharper than the original fused data (Fig.5e). The gain in resolution is made possible thanks to the recovery of an accurate PSF estimation, shown in Fig. 5d. We notice that the recovered PSF is elongated towards orthogonal directions, as expected in the reconstructions based on multi-view LSFM. We have previously estimated the elongation of the point spread function along the scanning axis (the worse resolution in the image); thus, we compare it with the profile along the elongated direction of the recovered PSF. The resulting graph is plotted in Fig. 5f. Our method blindly recovers a PSF close to the measured one, guarantying a resolution increase even when completely ignoring the optical response of the system.

**Figure 5.**
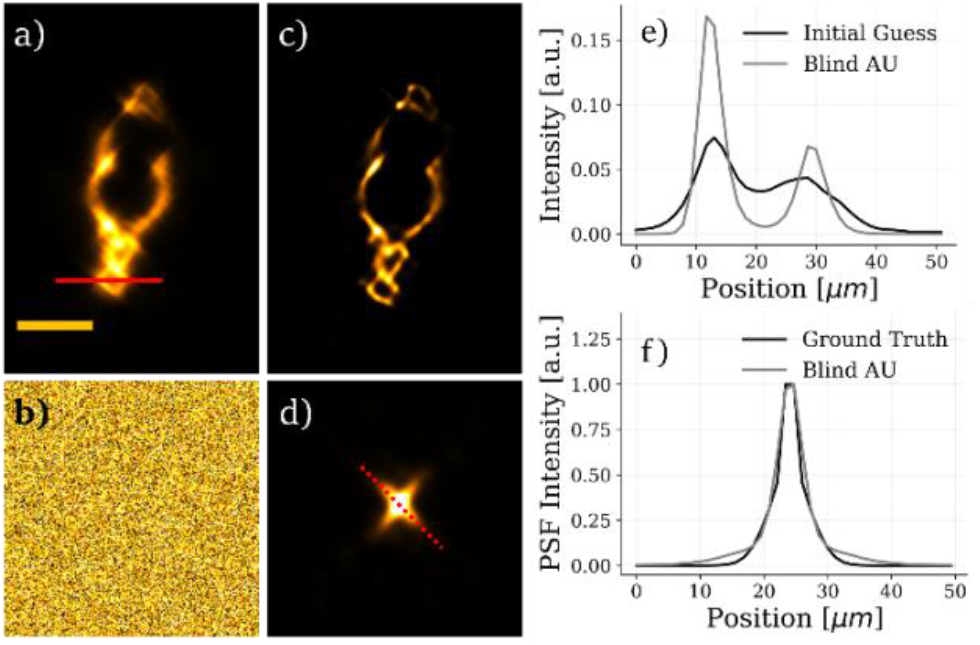
Multi-view Blind-AU reconstruction of a sample slice starting with a random PSF initial guess. Blind-AU algorithm applied on a tomographic selected slice of the four orthogonal views of the tail of a zebrafish embryo (45°, 135°, 225°, 315°). (a) Initial guess for the sample, corresponding to the aligned fusion of the four orthogonal views. (b) Random initial guess for the PSF. We assume to have null information about the structure of the PSF of our volume. (c) Tomographic slice reconstructed with our Blind-AU method. (d) The PSF obtained after the blind optimization routine. (e) Intensity profiles plotted along the red line traced in (a) for the sample initial guess and Blind-AU reconstruction. (f) Intensity profiles plotted along the red dashed line in (d) for the PSF ground truth and Blind-AU reconstruction.

## Conclusions

In this article, we have introduced a new method to form aligned reconstructions in multi-view light-sheet microscopy. Compared to the state-of-the-art algorithms, our implementation obtains sharp and inherently aligned reconstructions in the case where we have limited knowledge of the optical response of the measuring setup. To achieve this, we have generalized the AU algorithm [13], which we used for alignment-free multi-view reconstruction imaging [14], by including the blind deconvolution feature. Our choice was to alternate the reconstruction of the object with that of the point-spread function, keeping one fixed while solving for the other. To this end, we took inspiration from blind strategies grounded on the Richardson-Lucy deconvolution approach [15],[17] but here, instead, starting from the estimation of a global autocorrelation. We have already proven that autocorrelations can form implicitly aligned reconstructions down to the sub-pixel level [14]; thus, we decided to extend further its range of applicability. Thanks to its design, our method forms accurate reconstructions from multi-view acquisitions without knowing the PSF of the system. The validation was assessed under three different regimes and followed the actual software development. Initially, a blurred synthetic specimen was reconstructed by fusing multi-views in the autocorrelation domain, proving the fidelity of the reconstruction against the known ground truth. Once validated numerically, we tested it with experimental acquisitions of a three-dimensional specimen. We examined the vasculature of a zebrafish, a well-known model organism that permits us to spot the formation of eventual artifacts in the reconstructions. These reconstructions were compared with those obtained by standard deconvolution routines, providing contrast increase. In the ultimate analysis, we have assessed the correct recovery of the PSF in our multi-view LSFM setup, comparing it with the measured optical response along the scanning axis. The setup used in our experiment was a custom-made light-sheet fluorescence microscope with a rotating stage but, our technique is readily usable in any omnidirectional LSFM setups [19], or in techniques directly based on autocorrelation inversion such as hidden tomography [20].

Although it guarantees excellent reconstruction capabilities, our method may have room for further improvements. The AU algorithm is, in fact, a slowly converging method [14] which we iterate blindly for tens to hundreds of iterations within each cycle. Experimenting with a different number of blind iterations and the careful choice of initial guesses may speed up the recovery process. Similar Bayesian approaches have already benefit enormous speed-up by choosing ad-hoc backpropagating kernels [9]. This idea is suitable for our implementation and can lead to faster solutions. Including priors could also enhance the performance of our method. For example, in multi-view geometry, the angle of rotation is generally known. Designing a functional form for the PSF that accounts for the angular rotation could ease the reconstruction process. Lastly, our algorithm assumes uniform PSF blurring thorough the entire imaged volume. Although this is often considered a good approximation, higher image quality can be reached when considering spatially anisoplanatic deconvolutions [21] or mixed optical aberration corrections [22]. These considerations fall beyond the scope of the present manuscript, which may open new paths for further studies.

## Materials and methods

### Synthetic sample generation and

To test the Blind AU algorithm, we generated an image simulating blood vessels. The simulated multi-view measurement is composed of two orthogonal views of the sample, generated by convolving the original image with an elongated Gaussian PSF (with standard deviation *σ*_*z*_ = 2.7 px, *σ*_*x*_= 1 px), oriented along two different axes (*θ*_1_ = 0° and *θ*_2_ = 90°). Then, images are normalized to 2^16^ and Poisson noise is added, with average and variance equal to *λ* = 2^8^. For N different views, identified by the subscript *θ*_*i*_, each simulated measurement is determined by:

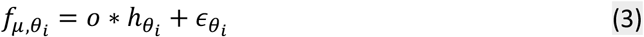

Where 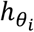 is the PSF rotated by the angle *θ*_*i*_, and 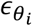 is the Poisson noise added to every single view. Each matrix has an odd dimension of 191×191 pixels, so the rotation of the PSF is applied around the central pixel, and the Gaussian is perfectly symmetric with respect to the center of the image. The resulting *f*_*μ*,1_ and *f*_*μ*,2_ are displayed in Fig. 2.

### Light sheet fluorescence microscopy

Light-sheet fluorescence microscopy is an optical technique that exploits decoupled illumination and detection, placed orthogonal to each other, to achieve optical sectioning. The illumination arm generates a light sheet (a single plane of illumination) on the sample. The emitted fluorescence is collected along the detection path, filtered, forming an image on a widefield detector (CMOS camera). Having independent illumination and detection, the PSF is determined by the product between illumination and detection PSFs and is elongated, typically, along the axial direction [23]. The system starts with a single-mode blue laser emitting at 473 *nm* (MBL-FN-473, 50 *m*W power), operating in CW mode. The laser beam is collimated to a diameter of 4 *mm* (Thorlabs collimator RC04FC-P01), and its width is reduced to 1 *mm* by a vertical slit. Then, the light is sent on a galvanometer (Thorlabs GVS001) conjugated with the sample position along the optical illumination path. The galvanometric mirror is controlled by a sinusoidal voltage signal that oscillates with an amplitude of ± 1° and frequency of 250 *Hz*. This pivots the illumination beam to remove shadowing artifacts that are common for side illumination of the sample [24]. Then, the beam is expanded by a factor of 3 by a telescope (4f system, with *f*_1_ = 50 mm and *f*_2_ = 150 *mm*), and a cylindrical lens (*f* = 200 *mm*) generates the light sheet, focusing light only along the vertical direction. The beam diameter is halved by a second 4f system (*f*_1_ = 400 *mm*, *f*_2_ = 200 *mm*), and light is collimated towards the back focal plane of the illumination objective (Olympus water immersion objective, 10x, *NA* = 0.3). The emitted fluorescence is collected by a second identical objective, mounted in a perpendicular direction with respect to the illumination path. Then, light crosses a GFP filter (center wavelength of 520 nm), the tube lens (*f* = 180 *mm*), and is collected by a low noise sCMOS camera (Neo 5.5 sCMOS Andor Technology). The size of the beam after the vertical slit results in an illumination numerical aperture *NA*_*ill*_ = 0.0055 and an axial resolution *δz*_*ill*_ = 5.65 *μm*, whereas the lateral resolution is *δρ*_*det*_ = 0.87 *μm*. In the central unit of the microscope, the sample is mounted fixed to the mechanical support that controls its position and orientation using a translator (PI M-403) and a rotator (PI M-037). Volumes are acquired by scanning the sample through the illumination beam, with minimum exposure time for each slice (0.0101305 *s*) and displacement along the detection axis between two consecutive acquisitions of 1.3 *μm*. This value is also the lateral dimension of the region imaged by every single pixel, by applying a 2 × 2 binning to the camera.

### PSF characterization

The point-spread function (PSF), representing the intensity impulse response of the microscope, describes the image degradation process. The PSF was estimated experimentally by measuring a sample composed of sparse fluorescent nanobeads, much smaller than the microscope resolution: they act as point sources and are transferred to the image plane according to the impulse response of the system. Beads should be measured under the same experimental conditions as the sample, avoid variations due to wavelength changes, and retrieve information about the alignment of the optical elements [25]. We used fluorescent nanobeads (Estapor, XC dye, 100 nm size) absorbing between 460 *nm* and 500 *nm* and emitting in the 515 – 570 *nm* range. They are embedded in phytogel with a concentration of 1: 105 and inserted in a cylindrical FEP tube with an internal diameter of 2 *mm* and an external one of 4 *mm*. To characterize the 3D PSF, we acquired a full volume of beads at the same experimental conditions as the sample; then, we selected a fluorescent bead in the center of the FOV and applied a Gaussian fit along the three dimensions to retrieve an analytical expression.

### Zebrafish handling and preparation

Adult zebrafish were maintained according to National (Italian D.lgs 26/2014) and European laws (2010/63/EU and 86/609/EEC), controlling experiments on live animals. Embryos, collected by natural spawning, were staged and raised at 28°C in fish water (Instant Ocean, 0.1% methylene blue). We used zebrafish embryos (*Danio rerio*) belonging to the line Tg(fli1a:EGFP)y1. The entire vascular tree can be visualized in vivo, thanks to the expression of Enhanced Green Fluorescent Protein (EGFP) under the control of the endothelial-specific gene promoter fli1a [26],[27]. At 48 hours post-fertilization, embryos have a typical dimension of 700 × 700 × 3000 *μm*^3^. They are anesthetized and mounted in Fluorinated Ethylene Propylene (FEP) tubes (outer diameter of 1.6 mm, inner diameter of 0.8 *mm*, 3 *cm* long), filled with 0.1% agarose, and a final tricaine concentration of 160 *mg*/*l* [28]. The tube is mounted in the microscope chamber, filled with a stock solution composed of fish water with 0.016% tricaine concentration.

### Richardson Lucy deconvolution

Richardson-Lucy deconvolution (RLD) algorithm is an iterative numerical method for image restoration, stable in presence of high noise levels [15],[16]. It is derived by interpreting images and PSFs as probabilities and applying Bayes theorem. In microscopy, the system’s point spread function, *h*, is not ideal; thus, the measurement *f*_*μ*_ of an object *o* can be described by the convolution *f*_*μ*_ = *o* ∗ *h* + *ϵ*. In this description, we can state that pixel-value of *f*_*μ*_ is determined by the re-distribution of the energy of *o* accordingly to *h*. Assuming that the PSF is shift-invariant (isoplanatic condition), the RLD iterative process can be written in a convolutional form:

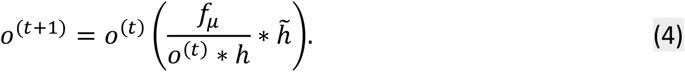

With the notation 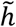, we indicate the PSF with reversed coordinates: 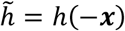, and the index *t* identifies iterations. Starting from an initial guess *o*^0^ ≥ 0, nonnegativity is preserved during the iteration process, without the need for positivity constraints [17]. Even if not explicitly reported, we monitored the reconstruction quality with the signal-to-noise ratio (SNR), expressed in decibel (dB), applied to the generic signal as:

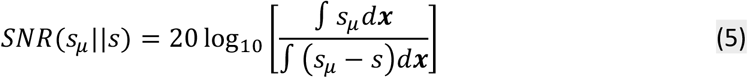

Where *s*_*μ*_ is the measurement of the signal (in the case of images, it is blurred and corrupted by noise), and *s* is the noiseless estimation of the signal measurement at a certain iteration.

### Pre-processing of experimental data

In the present manuscript, a multi-view measurement is composed of four orthogonal views, acquired by rotating the sample by steps of 90°. First, we subtracted the background signal from each matrix by measuring a full stack without laser illumination. Second, we rotate each stack against the reference view: the passage is performed in multiples of 90°. The measure passed to the algorithm lies in the autocorrelation space, and it is computed by autocorrelating each stack and averaging them. It guarantees intrinsic sub-pixel alignment [14]. Here, autocorrelation is carried out as multiplication in the Fourier domain, according to the cross-correlation theorem. To avoid numerical errors before the fusion, we take the absolute value of each autocorrelation. To determine an initial guess for the sample image, we decided to align the views. This step is carried out by setting the first view as a reference, then computing the cross-correlation between this stack and the others. For each couple, the position of the cross-correlation maximum represents the translational shift to apply in the three spatial coordinates. Finally, the initial guess is computed as the average of all the aligned views. In addition, we removed all the null elements by substituting a small value, taken as 1/100 of the image average. Otherwise, those pixels will remain zero during deconvolution. It is worth noticing that the alignment procedure is not strictly necessary to run the algorithm but ensures convergence in case the object is truncated in the image plane.

## Acknowledgments

We are grateful to Giovanni Vitale, Luca Persani, Silvia Carra and Germano Gaudenzi (Istituto Auxologico Italiano, IRCCS) for providing the biological samples and for helping in the zebrafish preparation. This work was supported by Horizon 2020 (EU) Marie Skłodowska-Curie Actions (HI-PHRET project, 799230) and Laserlab Europe V (871124).

